# Telmisartan potentiates insulin secretion via ion channels, independent of the AT1 receptor and PPARγ

**DOI:** 10.1101/2020.09.20.305334

**Authors:** Tao Liu, Lijuan Cui, Huan Xue, Linping Zhi, Xiaohua Yang, Huanhuan Yang, Mengmeng Liu, Tao Bai, Zhihong Liu, Min Zhang, Qing Guo, Peifeng He, Yunfeng Liu, Yi Zhang

## Abstract

Angiotensin II type 1 receptor blockers (ARBs), as antihypertensive drugs, have drawn attention for their benefits to individuals with diabetes and prediabetes. However, the effects of ARBs on insulin secretion remain unclear. Here, we investigated the insulinotropic effects of ARBs (telmisartan, valsartan, and irbesartan) and the underlying electrophysiological mechanism in rat islets. We found that only telmisartan among the three ARBs exhibited an insulin secretagogue role. Distinct from other ARBs, telmisartan exerted effects on ion channels including voltage-gated potassium (Kv) channels and voltage-gated Ca^2+^ channels to promote extracellular Ca^2+^ influx, thereby potentiating insulin secretion in a glucose-dependent manner. We observed that the peroxisome proliferator-activated receptor γ pathway was not involved in these telmisartan-induced effects. Furthermore, we identified that telmisartan at least directly inhibited Kv2.1 channel through construction of a Chinese hamster ovary cell line with Kv2.1 channel overexpression. Acute exposure of type 2 diabetes model (*db*/*db*) mice to a telmisartan dose equivalent to therapeutic doses in humans resulted in lower blood glucose and increased plasma insulin concentration in the oral glucose tolerance test. We further observed the telmisartan-induced insulinotropic and electrophysiological effects on pathological pancreatic islets isolated from *db*/*db* mice. Collectively, our results establish an important function of telmisartan distinct from other ARBs in the treatment of diabetes.

## Introduction

Diabetes and hypertension constitute common clinical conditions that are interlinked through numerous pathophysiological mechanisms *(Deedwania, 2004; Ferrannini and Cushman, 2012)*. In particular, hypertension substantively increases the risk of type 2 diabetes mellitus (T2DM), as revealed by a prospective cohort study wherein subjects with hypertension were almost 2.5 times more likely to develop T2DM than those with normal blood pressure *(Gress T et al., 2000)*. In turn, the majority (70%–80%) of patients with T2DM also have hypertension *(Fox et al., 2015)*.The coexistence of both conditions significantly increases the risks of developing nephropathy, heart failure, and other cardiovascular disease, leading to high rates of mortality and morbidity *(Deedwania, 2004; Ferrannini and Cushman, 2012)*. Therefore, the identification of drugs that prevent both conditions would be of considerable clinical importance.

Growing evidences indicated that angiotensin II type 1 (AT1) receptor blockers (ARBs), an important drug class in the treatment of hypertension and heart failure, provided beneficial effects for patients with diabetes and prediabetes. Several clinical trials and retrospective-analyses have shown that ARBs reduce the incidence of new-onset diabetes among patients with hypertension and heart failure *(NAVIGATOR Study Group et al., 2010; Yusuf et al., 2005; Kjeldsen et al., 2006)*. Moreover, it has been repeatedly demonstrated that ARBs ameliorate T2DM and its related complications such as atherosclerosis and nephropathy *(Candido et al., 2004; Makino et al., 2008; Viberti et al., 2002; Parving et al., 2001)*. In addition, ARBs have been highly recommended in pharmacological therapy regimens for patients with both diabetes and hypertension by the American Diabetes Association *(American Diabetes Association, 2015)*.

T2DM is a metabolic disorder syndrome characterized by insulin resistance and deficiency. The confirmed benefits of ARBs in patients with diabetes and prediabetes have been primarily attributed to blockade of the local renin–angiotensin system (RAS). ARBs suppress oxidative stress and inflammatory responses resulting from overactivity of this system, thereby protecting β-cells against dysfunction and improving insulin sensitivity to maintain euglycemia *(van der Zijl et al., 2011; Hunyady and Catt, 2006; Li et al., 2012; Nagel et al., 2006; Shiuchi et al., 2004)*. However, although insufficient insulin secretion constitutes a fundamental process that determines the onset and progression of T2DM *(Weyer et al., 1999; Levy et al., 1998)*, few studies have focused on the effect of ARBs on insulin secretion or its underlying mechanism.

In the present study, we applied three ARBs, namely telmisartan, valsartan, and irbesartan to evaluate the effects of ARBs on insulin secretion and investigate the underlying electrophysiological mechanism. Notably, our data showed that unlike other ARBs, telmisartan glucose-dependently elevated the intracellular [Ca^2+^] ([Ca^2+^]_i_) levels of β-cells through its distinctive action on ion channels, leading to enhanced insulin secretion. Our findings suggest that in addition to the typical beneficial effects of ARBs, telmisartan may serve as an insulin secretagogue in the treatment of patients with both diabetes and hypertension.

## Results

### Telmisartan, but not valsartan or irbesartan, enhances glucose-stimulated insulin secretion (GSIS)

To examine the effects of ARBs on insulin secretion, firstly, isolated rat islets were treated with various doses of telmisartan. As shown in Fig. 1A, telmisartan (10 and 50 μM) potentiated insulin secretion under 8.3 mM glucose conditions but had no effect under 2.8 mM glucose. Furthermore, the data in Fig. 1 B confirmed that telmisartan-induced insulin secretion was glucose-dependent. Next, the functions of other ARBs were evaluated. Notably, no promotion of insulin secretion was observed following treatment with valsartan and irbesartan under 8.3 and 16.7 mM glucose conditions (Fig. 1, C and D, and fig. S1). Considering that telmisartan, valsartan and irbesartan are clinically available ARBs owing to their high specificity for AT1 receptors *(Michel et al., 2013)*, our results suggested that telmisartan-mediated insulinotropic effect was independent of AT1 receptors.

**Fig. 1:**
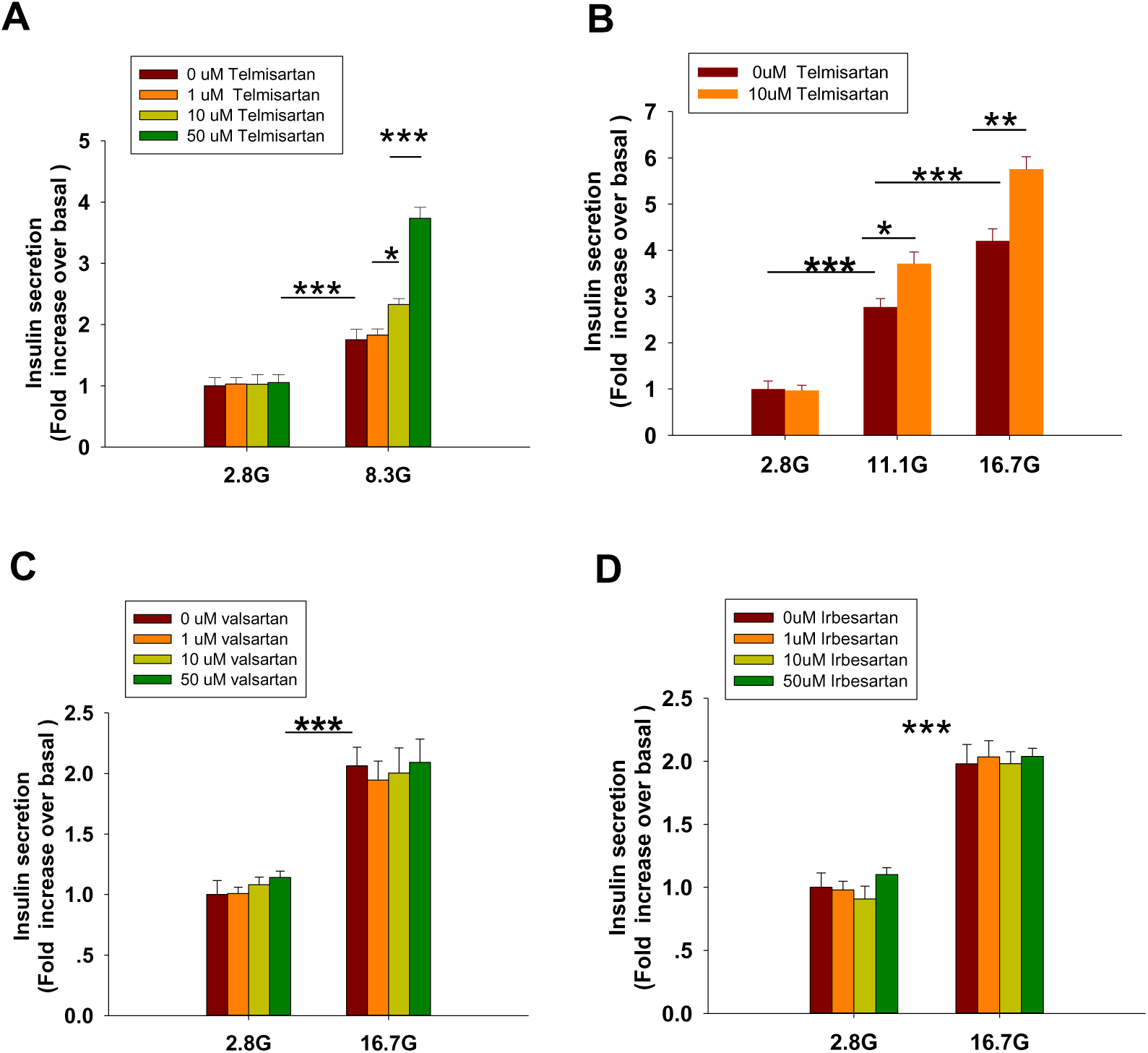
Only telmisartan among the three ARBs potentiated insulin secretion in rat islets. In every tube, five handpicked rat islets were incubated for 30 min in 500 μL Krebs Ringer bicarbonate-HEPES (KRBH) buffer containing different drugs and glucose concentrations, then supernatant liquid was collected for insulin measurement (n = 7 tubes per group). (A) Rat islets were treated with various doses (1, 10, and 50 μM) of telmisartan under 2.8 and 8.3 mM glucose (denoted as 2.8 G and 8.3 G) conditions. (B) Islets were treated with 10 μM telmisartan under different glucose concentrations (2.8, 11.1, and 16.7 mM). (C and D) Rat islets were treated with various doses (1, 10, and 50 μM) of valsartan or irbesartan under 2.8 and 16.7 mM glucose conditions. All results are normalized to basal secretion at 2.8 G, and reported as the means ± SEM. Statistical differences among three or more groups were compared using one-way analysis of variance (ANOVA) and Student–Newman–Keuls method post hoc analysis. Statistical differences between two groups (with or without telmisartan) under the same glucose condition in (B) were determined using an unpaired two-tailed Student’s *t* test. * P < 0.05, ** P < 0.01, *** P < 0.001.

### Telmisartan, but not valsartan or irbesartan, increases ([Ca^2+^]_i_) concentration in β-cells

Within β-cells, eliciting an increase in [Ca^2+^]_i_ causes insulin granule exocytosis; therefore, the elevation in [Ca^2+^]_i_ level is essential to induce insulin secretion *(Sabatini et al., 2019)*. To verify whether the insulinotropic effect of telmisartan was related to the change in [Ca^2+^]_i_, we applied the calcium-sensitive dye Fura 2-AM to detect changes in fluorescence intensity. Telmisartan (10 and 50 μM) induced an acute increase in fluorescence intensity dose-dependently. Moreover, the elevation only occurred under high (11.1 and 16.7 mM) (Fig. 2 C, D, and E, F) but not low (2.8 mM) (Fig. 2 A and B) glucose conditions. In addition, in the calcium imaging experiment, neither valsartan (Fig. 2 G and H) nor irbesartan (Fig. 2 I and J) increased the [Ca^2+^]_i_ concentration of β-cells under high glucose conditions (16.7 mM).

**Fig. 2:**
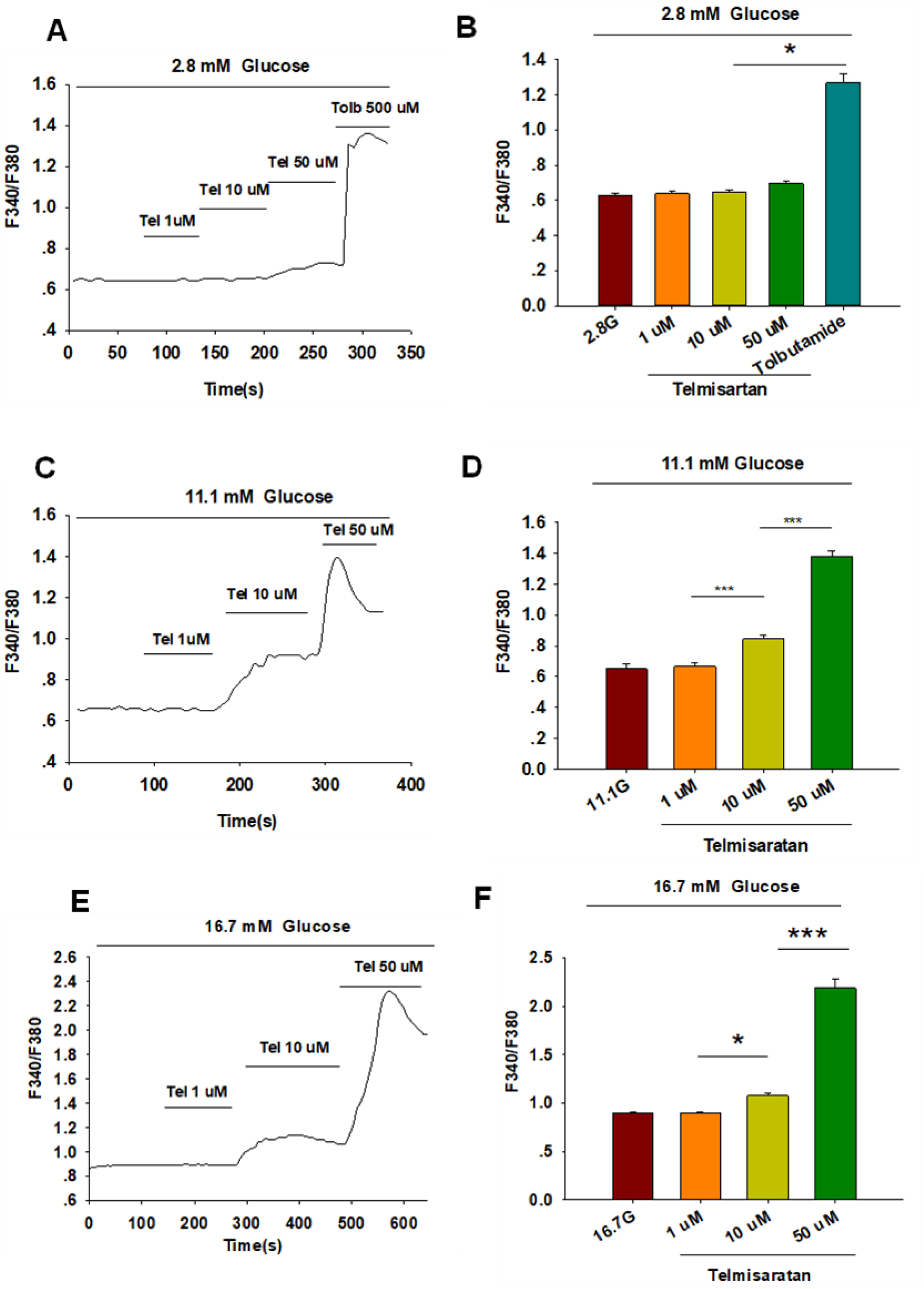
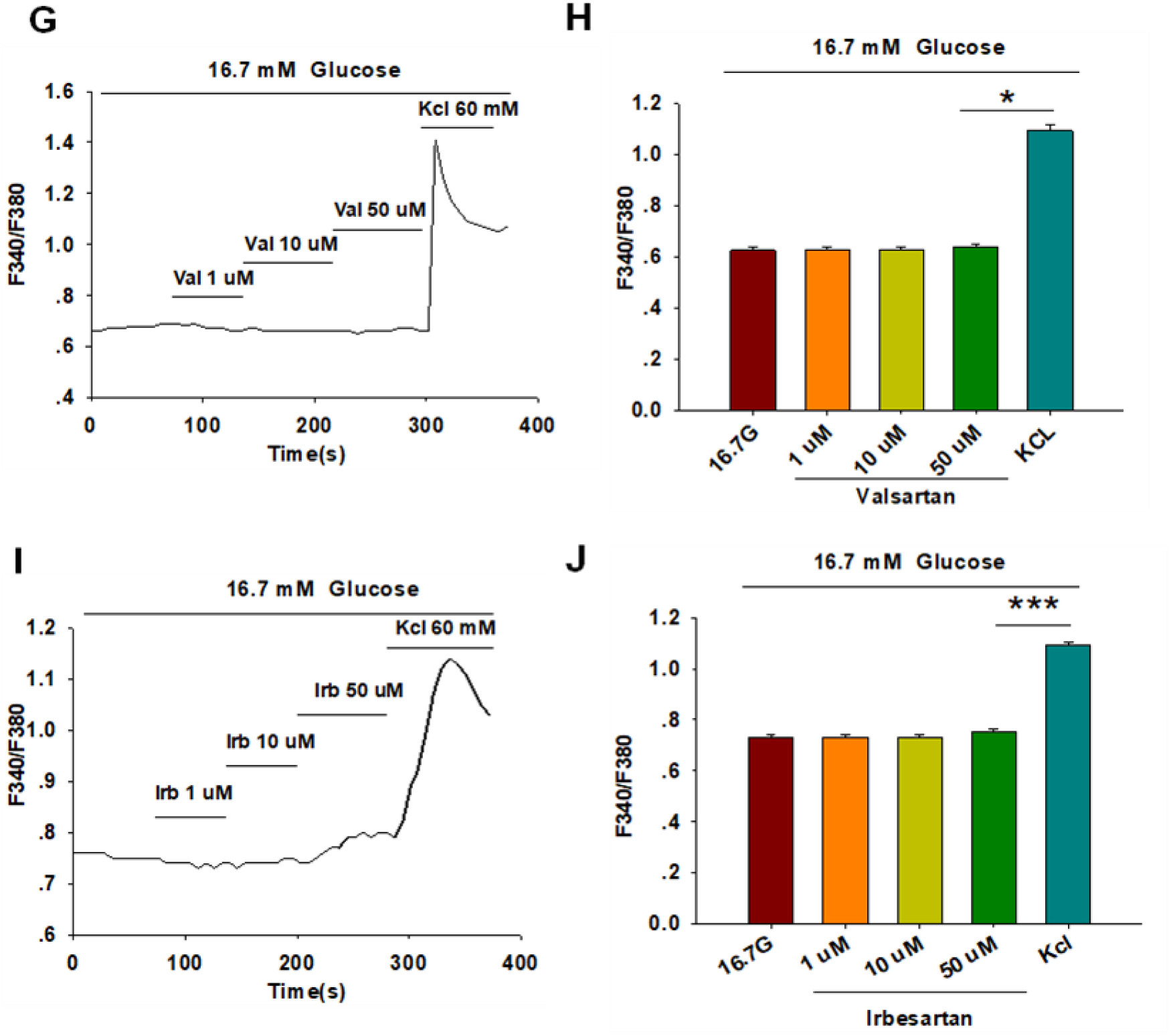
Only telmisartan among the three ARBs increased intracellular [Ca^2+^] ([Ca^2+^]_i_) concentration in rat pancreatic β-cells. (A) The trace shows the changes of [Ca^2+^]_i_ concentration in β-cells treated with 1, 10, and 50 μM telmisartan (Tel) under 2.8 mM glucose conditions; 500 μM tolbutamide (Tolb) was used as a positive control. (B) The average value during 30 s F340/F380 spikes for each test in response to different doses of telmisartan under 2.8 mM glucose conditions as indicated (n = 9). (C and D) The trace shows the changes of [Ca^2+^]_i_ concentration in β-cells treated with different doses of telmisartan under 11.1 mM glucose conditions, and the average value during 30s F340/F380 spikes for each test as indicated (n = 9). (E and F) The trace shows the changes of [Ca^2+^]_i_ concentration in β-cells treated with different doses of telmisartan under 16.7 mM glucose conditions, and the average value during 30s F340/F380 spikes for each test as indicated (n = 9). (G, H and I, J) The trace shows the changes of [Ca^2+^]i concentration in β-cells treated with 1, 10, and 50 μM of valsartan (Val) or irbesartan (Irb) under 16.7 mM glucose conditions respectively, and the average value during 30s F340/F380 spikes for each test as indicated. KCl (60 mM) was used as a positive control (n = 9). All results are reported as the means ± SEM. Statistical differences among three or more groups were compared using one-way ANOVA, and followed by Student– Newman–Keuls Method post hoc analysis in (D), (F), and (J), or Tukey post hoc analysis in (B) and (H). * P < 0.05, *** P < 0.001

### Peroxisome proliferator-activated receptor γ (PPARγ) is not involved in telmisartan-induced insulin secretion and elevation of [Ca^2+^]_i_ levels

Telmisartan and irbesartan have also been reported to function as a partial agonist of PPARγ *(Schupp et al., 2004, 2005)*. In view of the absence of changes in insulin secretion and [Ca^2+^]_i_ levels with irbesartan, we speculated that PPARγ might not be responsible for the effects of telmisartan on these measures. We therefore performed the insulin secretion assay and calcium imaging experiment using GW9662, a selective PPARγ antagonist. As shown in Fig. 3, GW9662 alone had no effect on GSIS and [Ca^2+^]_i_ concentration, and the addition of GW9662 did not influence the effects of telmisartan on insulin secretion or [Ca^2+^]_i_ concentration.

**Fig. 3:**
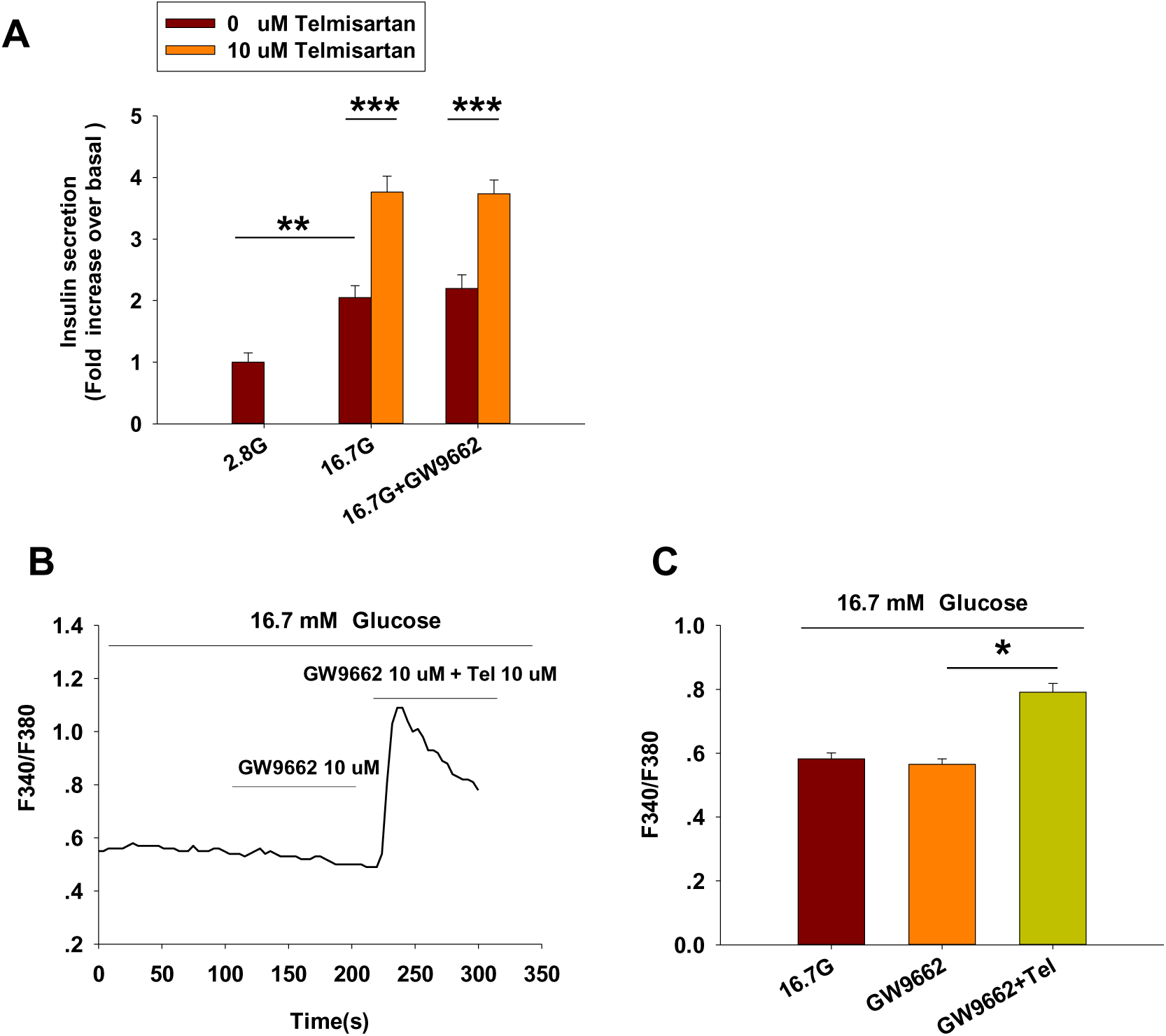
PPARγ does not participate in the pathway of telmisartan-induced insulin secretion and elevation of [Ca^2+^]_i_ levels (A) Telmisartan (10 μM) potentiated glucose-stimulated insulin secretion in the presence or absence of the PPARγ inhibitor GW9662 (10 μM) (n = 7). All insulin secretion results are normalized to basal secretion at 2.8 Mm glucose concentration. (B) The trace shows the changes of [Ca^2+^]_i_ concentration in β-cells treated with GW9662 (10 μM) alone or in combination with telmisartan (Tel 10 μM) under 16.7 mM glucose conditions. (C) The average value during 30s F340/F380 spikes for each test in response to GW9662 alone or in combination with telmisartan under 16.7 mM glucose conditions as indicated (n = 9). All results are reported as the means ± SEM. In (A), statistical differences between two groups (with or without telmisartan) were compared using an unpaired two-tailed Student’s *t* test, and difference among three groups without telmisartan were compared using one-way ANOVA and Student-Newman-Keuls method post hoc analysis. In (C), difference among three groups was determined by one-way ANOVA and Tukey Test post hoc analysis. *P < 0.05, **P < 0.01, ***P < 0.001.

### Telmisartan affects [Ca^2+^]_i_ concentration through extracellular Ca^2+^ influx rather than intracellular Ca^2+^ stores release

[Ca^2+^]_i_ levels of β-cells are tightly maintained through the regulation of extracellular Ca^2+^ influx and the movement of Ca^2+^ within intracellular stores *(Sabatini et al., 2019)*. We thus examined the concentration of [Ca^2+^]_i_ in the absence of extracellular Ca^2+^ to study the pathway by which telmisartan increases [Ca^2+^]_i_ concentration. Calcium imaging (Fig. 4 A and B) showed that telmisartan-induced elevation of [Ca^2+^]_i_ levels was reversed in Ca^2+^-free KRBH medium, although the [Ca^2+^]_i_ level was considerably increased upon intracellular Ca^2+^ mobilization via thapsigargin.

**Fig. 4:**
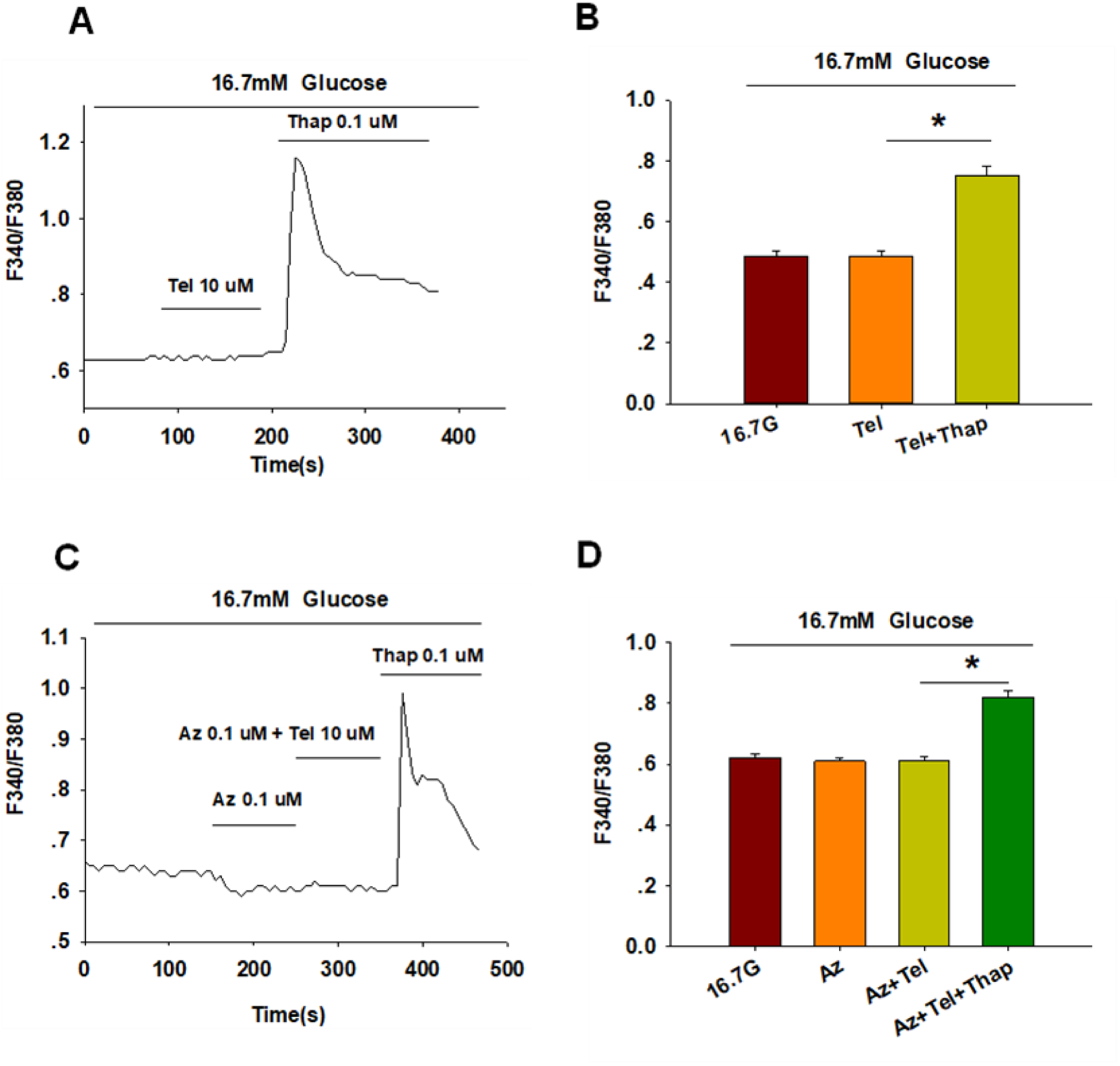
Telmisartan enhances [Ca^2+^]_i_ levels through extracellular Ca^2+^ influx, rather than intracellular Ca^2+^ stores release. (A) The trace shows the changes of [Ca^2+^]_i_ concentration in β-cells treated with telmisartan (Tel, 10 μM) under 16.7 mM glucose conditions in Ca^2+^-free KRBH medium. (B) The average value of F340/F380 during each test in response to telmisartan (Tel, 10 μM) in Ca^2+^-free KRB medium (n = 23). (C) The trace shows the changes of [Ca^2+^]_i_ concentration in β-cells treated with telmisartan (Tel 10 μM) under 16.7 mM glucose conditions with addition of the L-type VGCC blocker azelnidipine (Az, 0.1 μM). (D)The mean value of F340/F380 during each test in response to telmisartan (10 μM) with added azelnidipine (0.1 μM). Thapsigargin (Thap,0.1 μM) was used as a positive control (n = 12). All results are reported as the means ± SEM. Statistical differences among three or more groups were determined by one-way ANOVA, followed by Tukey Test post hoc analysis. * P < 0.05.

Moreover, telmisartan-induced effects on [Ca^2+^]_i_ levels were monitored following the application of azelnidipine, an L-type voltage-gated Ca^2+^channel (VGCC) blocker. We observed that the increase in [Ca^2+^]_i_ levels with telmisartan was completely blocked by azelnidipine, supporting that telmisartan enhances extracellular calcium influx through L-type VGCCs. Conversely, significant elevation remained upon thapsigargin addition (Fig. 4 C and D), confirming the lack of telmisartan effect on intracellular calcium stores.

### Telmisartan inhibits voltage-gated potassium (Kv) channels, and prolongs action potential durations (APDs) in β-cells

Pancreatic β-cells are electrically excitatory. Previous studies have demonstrated that Kv channels play an important role in GSIS and glucose-stimulated increase of [Ca^2+^]_i_ *(Herrington et al., 2006; Roe et al., 1996; MacDonald and Wheeler, 2003)*; therefore, we applied patch-clamp techniques to explore the effects of telmisartan on the Kv channels of β-cells. Fig 5 A and B illustrate that telmisartan decreased the Kv channel currents compared with that of controls.

**Fig. 5:**
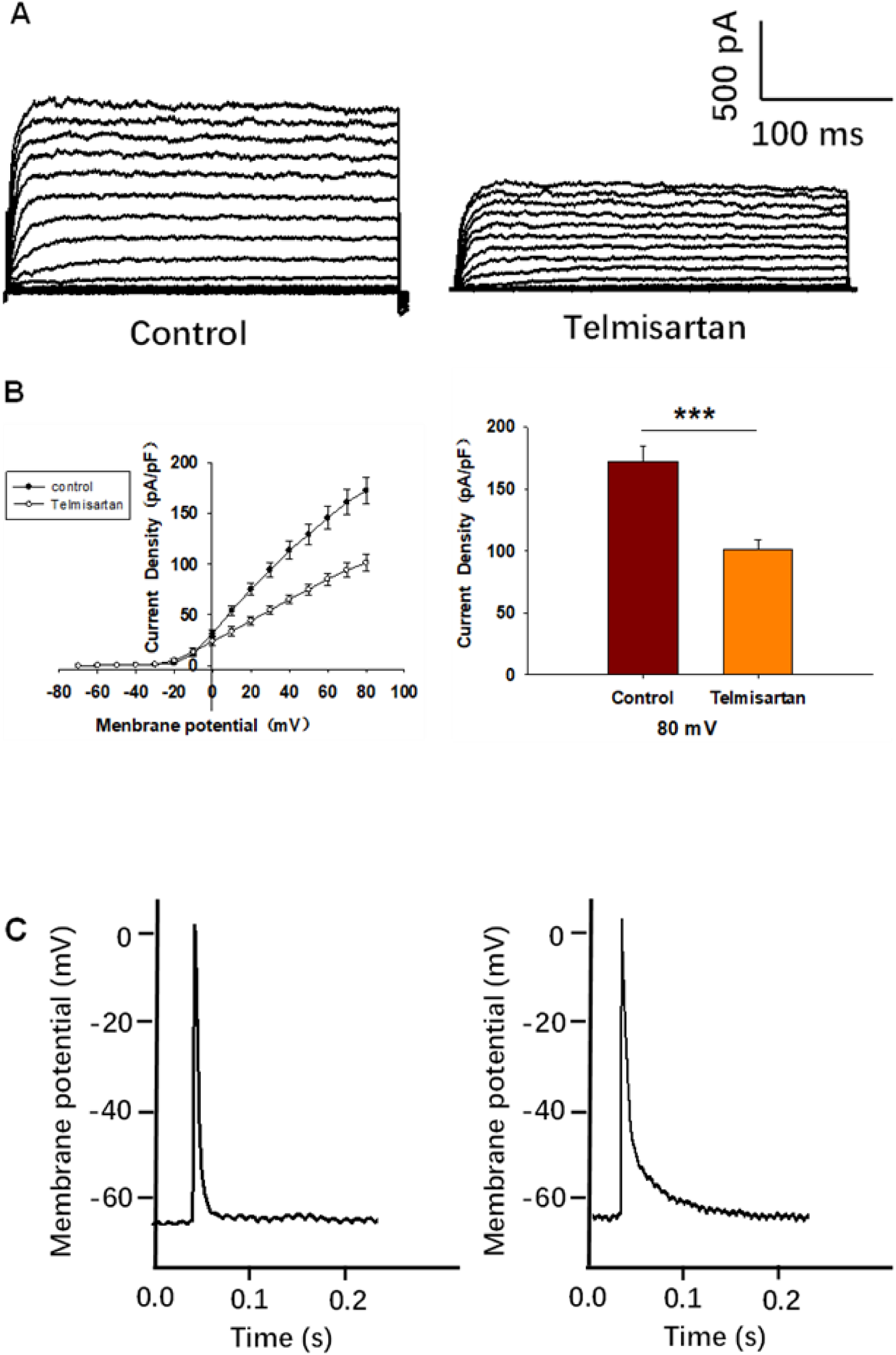
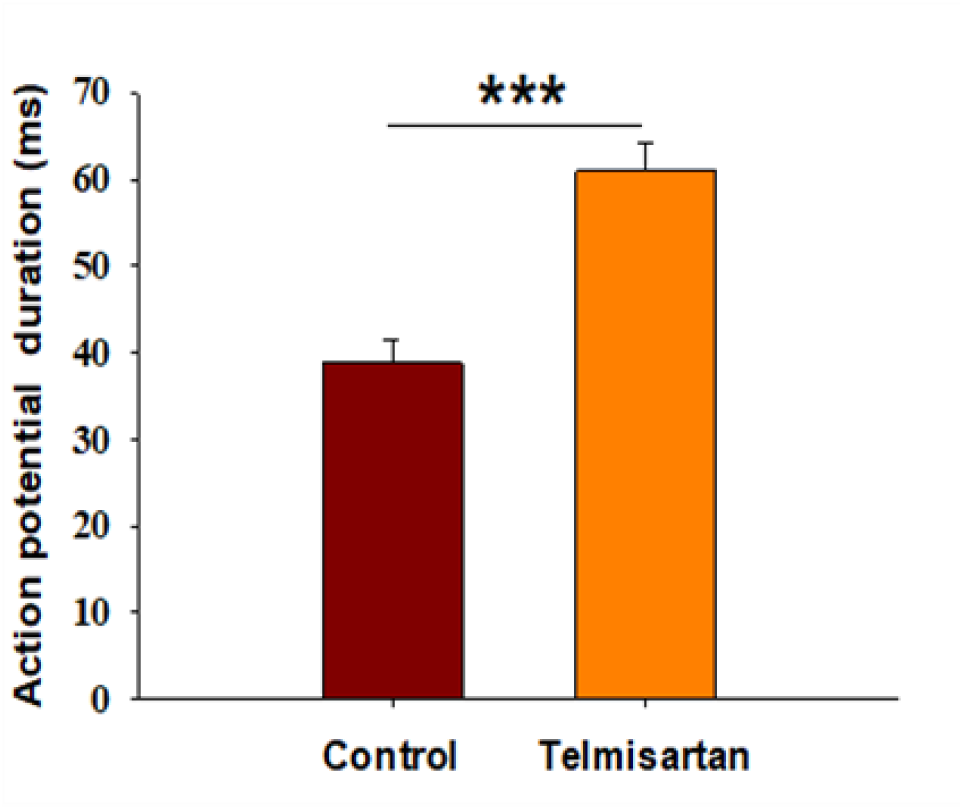
Pancreatic β-cells treated with telmisartan exhibit reduced Kv currents and extended APD. (A) Kv currents were recorded in voltage-clamp mode with holding potential from −70 to +80 mV in 10 mV increments. Representative current traces recorded in control and treated with telmisartan-treated (10 μM) β-cells. (B) Current-voltage relationship curves of Kv channels and summary of the mean current density of Kv channels recorded at 80 mV depolarization (control n = 9, telmisartan n = 7). (C) Action potentials were elicited by 4 ms, 150 pA current. Representative action potential waveforms for β-cells treated without or with telmisartan (10 μM) and summary of the mean APDs (n = 7). Statistical differences between two groups were determined using an unpaired two-tailed Student’s *t* test. *** P <0 .001.

Kv channels participate in the repolarization of action potentials of β-cells, so that inhibition of Kv channels delays the repolarization, thus prolonging the APD, namely the duration of extracellular Ca^2+^ influx *(Herrington et al., 2006; MacDonald and Wheeler, 2003; Jacobson and Philipson, 2007)*. Therefore, we next recorded the action potentials in current-clamp mode to observe the effect of telmisartan on APD. As presented in Fig. 5 C, comparison of APDs with or without telmisartan indicated that telmisartan extended APD.

### Telmisartan directly inhibits Kv2.1 channels independent of AT1 receptor and PPARγ

We evaluated the effects of valsartan and irbesartan in the voltage-clamp experiment. Neither valsartan nor irbesartan exhibited similar effects on Kv channels as those observed with telmisartan treatment (Fig. 6 A and B). Moreover, GW9662 addition did not influence the telmisartan-induced inhibition of Kv channels (Fig. 6 C and D). The results indicated that telmisartan inhibited Kv channels independent of the AT1 receptor and PPARγ.

**Fig. 6:**
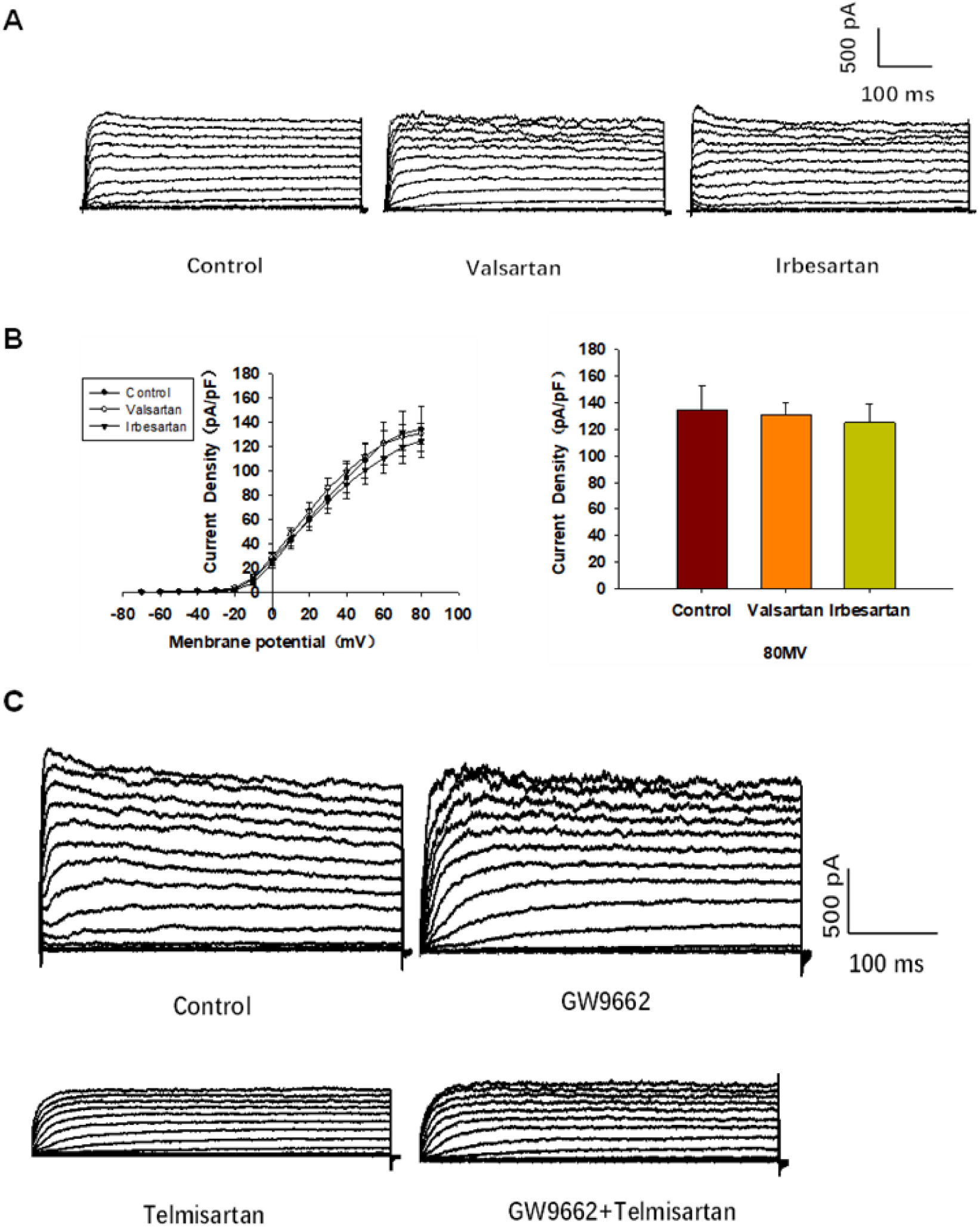
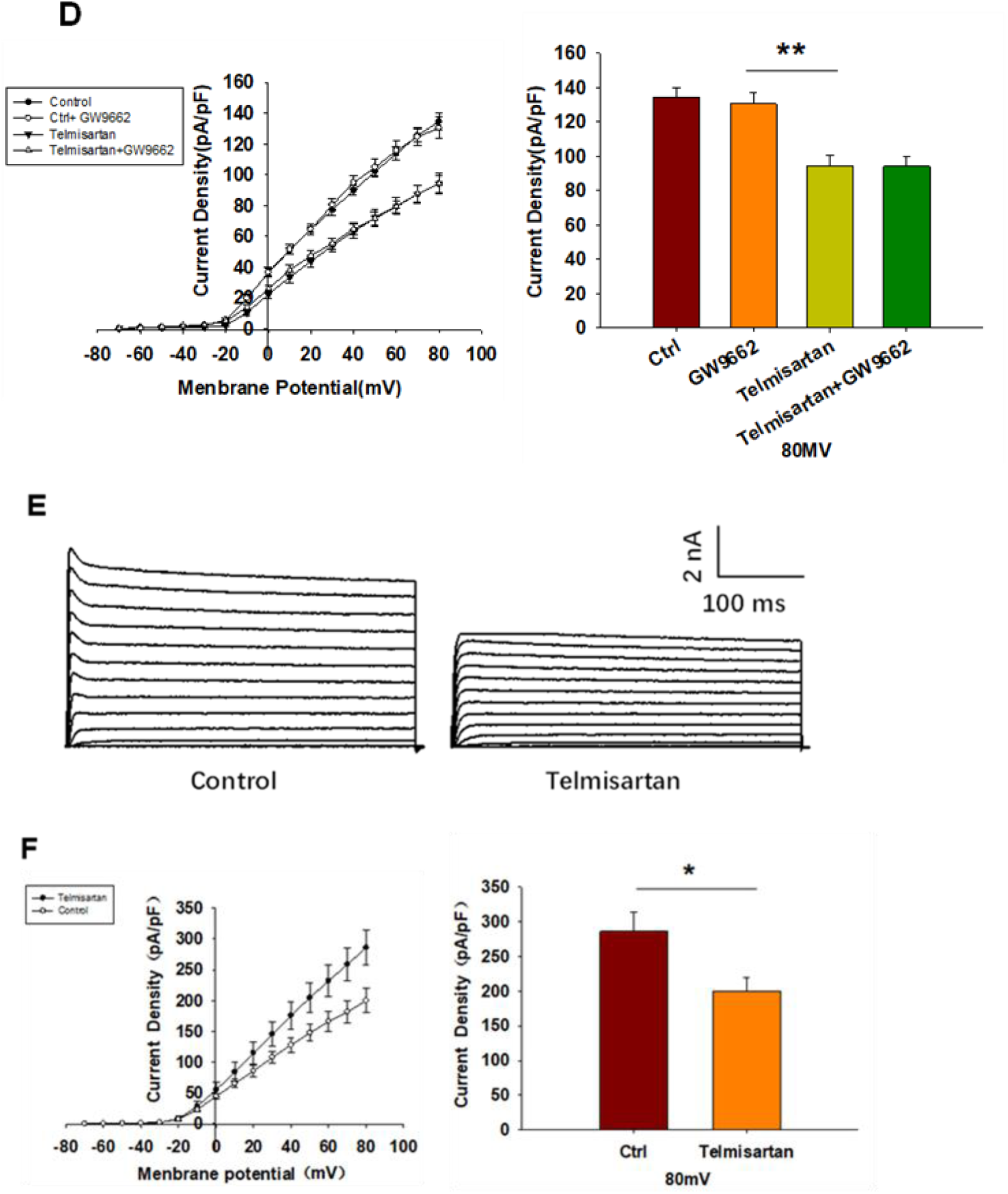
The AT-1 receptor and PPARγ are not involved in the telmisartan-induced inhibition of Kv channels, whereas telmisartan exerts a direct effect on Kv2.1 channels. (A) Representative current traces recorded upon treatment with valsartan (10 μM) and irbesartan (10 μM) in β-cells. (B) Current-voltage relationship curves and the summary of the mean current density of Kv channels recorded at 80 mV depolarization (control n = 7, valsartan n = 8, irbesartan n = 6). (C) Representative current traces recorded under treatment of telmisartan (10 μM) alone or in combination with GW9662 (10 μM) in β-cells. (D) Current-voltage relationship curves and the summary of the mean current density of Kv channels recorded at 80 mV depolarization (control n = 8, GW9662 n = 12, telmisartan n = 7, telmisartan+GW9662 n = 10). (E) The CHO-Kv2.1 cell line was constructed using a lentivirus vector overexpressing Kv2.1 channels. Representative current traces recorded without or with telmisartan (10 μM) in CHO-Kv2.1 cells. (F) Current-voltage relationship curves and the summary of the mean current density of Kv channels recorded at 80 mV depolarization (control n = 10, telmisartan n = 8). All results are reported as the means ± SEM. Statistical differences between two groups were determined using an unpaired two-tailed Student’s *t* test. Statistical differences among three or more groups were compared using one-way ANOVA. For comparing the effects of GW9662 groups, Tukey Test post hoc analysis was applied. * P < 0.05, ** P < 0.01.

We therefore hypothesized that telmisartan might directly inhibit Kv channels. As the Kv2.1 channel constitutes the main subtype among Kv families involved in the regulation of insulin release by β-cells *(MacDonald P et al., 2001; Li et al., 2013; Jacobson et al., 2007)*, we carried out patch-clamp experiments to determine whether telmisartan directly inhibited Kv2.1 channels. Chinese hamster ovary (CHO)cells, which do not express any endogenous Kv channels *(Yu and Kerchner, 1998)*, were utilized to establish the Kv2.1-overexpressing CHO-Kv2.1 cell line. Under whole-cell voltage-clamp mode, Kv2.1 channel currents and their suppression by telmisartan were both detected in CHO-Kv2.1 cells (Fig. 6 E, and F), suggesting that telmisartan exerted direct inhibition on Kv2.1 channels.

### Telmisartan activates VGCCs independent of the AT1 receptor and PPARγ

To further confirm whether Kv channels alone are involved in mediating telmisartan-induced insulin secretion and increase of [Ca^2+^]_i_ levels, tetraethylammonium chloride (TEA), a potent inhibitor of Kv channels, was employed in pancreatic β-cells. Previous studies have shown that 20 mM TEA blocks the majority of Kv channels and causes calcium elevation *(Roe et al., 1996; MacDonald P et al., 2001)*. As shown in Fig. 7A, TEA stimulated insulin secretion under 11.1 mM glucose conditions and telmisartan still significantly promoted insulin secretion in the presence of TEA, indicating that other factors may participate in telmisartan-stimulated insulin secretion. Consistent with this results, telmisartan also enhanced the [Ca^2+^]_i_ concentration in the presence of TEA under 11.1 mM glucose conditions (Fig. 7 B and C).

**Fig. 7:**
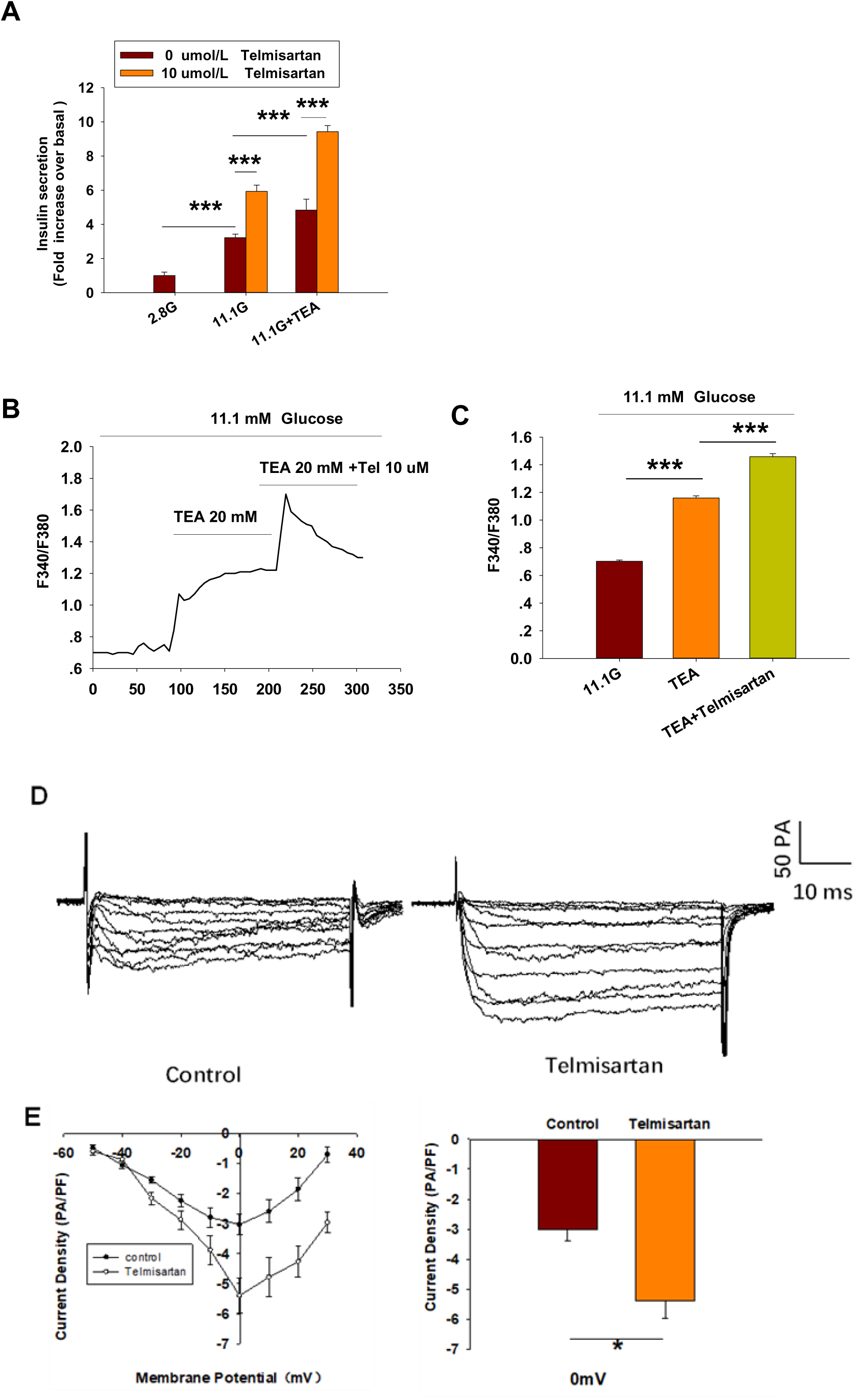
Kv channels partly mediate telmisartan-induced insulin secretion and increase of [Ca^2+^]_i_ levels. (A) Rat islets were treated with telmisartan (10 μM) in the presence or absence of TEA (20 mM) under 2.8 and 11.1 mM glucose conditions and insulin secretion was measured (n = 7). All insulin secretion results are normalized to basal secretion at 2.8 Mm glucose condition. (B) The trace shows the changes of ([Ca^2+^]_i_) concentration in β-cells treated with 20 mM TEA and in combination with 10 μM telmisartan (Tel) under 11.1 mM glucose conditions. (C) The average value during 30 s F340/F380 spikes for each test (n = 9). (D) VGCCs were recorded in voltage-clamp mode with test potentials from −50 to 30 mV in 10 mV increments. Representative current traces recorded in control and telmisartan-treated (10 μM) β-cells. (E) Current-voltage relationship curves of VGCCs and summary of the mean Ca^2+^current density recorded at 0 mV depolarization (control, n = 7; telmisartan, n = 8). All results are reported as the means ± SEM. Statistical differences between two groups were determined using an unpaired two-tailed Student’s *t* test. Statistical differences among three groups were compared using one-way ANOVA and Student–Newman–Keuls method post hoc analysis. Effects on VGCCs between telmisartan and control were compared using the Mann–Whitney Rank Sum Test. * P < 0.05, and *** P < 0.001.

As telmisartan enhances extracellular calcium influx through VGCCs, we performed patch-clamp experiments to observe the effects of telmisartan on VGCCs in pancreatic β-cells. As presented in Fig. 7 D and E, telmisartan increased voltage-dependent inward Ca^2+^ currents densities compared with those of controls. In addition, no significant difference was observed when VGCC currents were recorded following treatment with valsartan or irbesartan (fig. S2, A and B); telmisartan-induced activation was not inhibited by GW9662 co-administration (fig. S2, C and D). The results thus demonstrated that telmisartan also activated VGCCs of β-cells; moreover, neither the AT1 receptor nor the PPARγ mediated this effect.

### Telmisartan ameliorates hyperglycemia by increasing insulin secretion in vivo and amplifies GSIS in vitro in db/db mice

We applied db/db mice as T2DM model mice to determine whether telmisartan induced hypoglycemic effects in vivo. Male mice were administered with telmisartan (15 mg/kg) or vehicle once by gavage at the age of 8 and 11 weeks, then the oral glucose tolerance test (OGTT) was performed to observe the effects of telmisartan on glucose response.

In 8-week-old mice, blood glucose levels monitoring revealed that glucose clearance at 30 min and thereafter was improved significantly in telmisartan-treated mice, and noticeable difference was observed when the glycemic response was measured via the area under the curve (AUC) compared with that of control (Fig. 8 A). However, although the time of peak blood glucose was similarly advanced to 15 min in 11-week-old mice, it was not until 90 min and 120 min (approximately 1 h later than in 8-week-old mice) that the blood glucose values were markedly lower than those of controls. Additionally, the AUC results showed no significant difference between the groups (Fig. 8 B). Furthermore, the levels of plasma insulin in the telmisartan-treated group were considerably higher than those in the control group at 15, 30, and 60 min with the AUC differing significantly between the groups (Fig. 8 C). Therefore, the glucose-lowering effect of telmisartan was accompanied by the increase in the levels of plasma insulin, suggesting that the hypoglycemic effects of telmisartan were a result of increased insulin secretion. We speculated that the glucose-lowering effect of telmisartan was delayed and weakened in 11-week-old mice, possibly owing to the progression of insulin resistance and the deterioration of β-cell function in db/db mice.

**Fig. 8:**
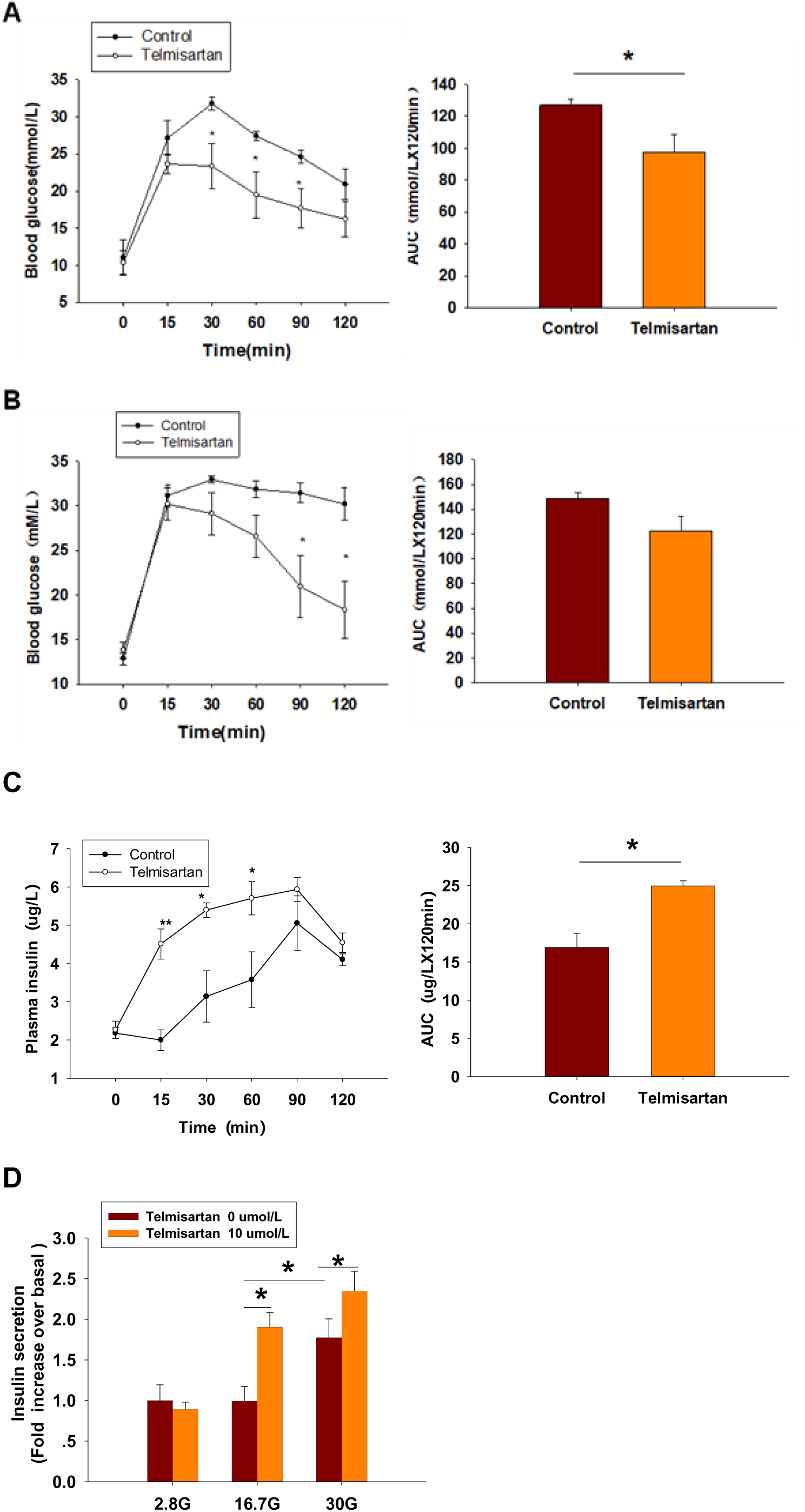
Telmisartan improves glucose tolerance in db/db mice, and elevates GSIS levels in isolated islets of db/db mice. After 14 h fasting, db/db mice were administered with telmisartan (15 mg/kg) or vehicle (0.5% carboxymethyl cellulose sodium) (n = 4 mice per group). After 2 h, mice were fed with glucose (1.5 g/kg), and glucose level and plasma insulin concentration in tail blood were determined. Finally, islets were isolated from the db/db mice to perform insulin secretion assays. (A) OGTT was performed and AUCs for OGTT were calculated from the data in 8-week-old mice. (B) OGTT and AUC for OGTT in 11-week-old mice. (C) Serum insulin levels at corresponding times and AUC in 11-week-old mice. (D) Db/db mice islets were treated with or without telmisartan (10 μM) under different glucose concentrations (2.8, 16.7, and 30 mM) (n = 6). All insulin secretion results are normalized to basal secretion at 2.8 Mm glucose concentration. All results are reported as the means ± SEM. Statistical differences between two groups were determined using the unpaired two-tailed Student’s *t* test unless otherwise stated. Glucose levels in 11-week-old mice at indicated time points were compared using the Mann–Whitney Rank Sum Test except at 0 min. AUCs calculated from the data of glucose levels or plasma insulin levels in 11-week-old mice were compared using the Mann–Whitney Rank Sum Test. As for the insulin assay in (D), statistical differences among three groups (without telmisartan) were compared using one-way ANOVA and followed by Student–Newman–Keuls method post hoc analysis, and differences between two groups under the same glucose conditions were compared using the paired *t* test. *P < 0.05, **P < 0.01.

At the end of the experiment, islets were isolated from *db*/*db* mice and used for an ex vivo study. The results showed that telmisartan potentiated insulin secretion under 16.7 and 30 mM glucose conditions (Fig. 8 D). Consistent with the results of in vivo studies, telmisartan treatment similarly enhanced GSIS under the pathological condition of diabetes. However, GSIS in cultured islets only occurred under 30 mM glucose conditions whereas 2.8 and 16.7 mM glucose showed equivalent secretion. The bluntness of GSIS might be related to impaired β-cell function caused by long-term exposure to high glucose and lipids in the development of diabetes in the db/db mice *(Olofsson et al., 2007)*.

In T2DM, high glucose and free fatty acids leads to adverse effects (including blunted GSIS and decreased cell viability) *(Olofsson et al., 2007; Tan et al., 2013)*, with the modulation of Kv and voltage-gated Ca^2+^ channels by “glucolipotoxicity” also being involved *(Hoppa et al., 2011; Lee et al., 2018)*. We next performed the patch-clamp experiments to ascertain whether telmisartan exerted similar electrophysiological effects on pathological β-cells. We observed that both the decreased Kv channel currents (fig. S3, A and B) and increased VGCC currents (fig. S3, C and D) remained in telmisartan-treated β-cells of db/db mice. Therefore, under the pathological condition of T2DM, telmisartan still served the function of an insulin secretagogue through its action on ion channels.

## Discussion

ARBs are of critical importance to individuals with both diabetes and hypertension. We therefore carried out the study to better understand the beneficial effects of ARBs for diabetes. Notably, we revealed an insulin secretagogue role for telmisartan, which is not present in other ARBs. In the present study, isolated islets were exposed to telmisartan for only 30 minutes prior to the insulin secretion assay, and glucose-lowering effects were observed in db/db mice following acute telmisartan administration. The evidences both in vitro and in vivo thus demonstrated the rapid insulinotropic effect of telmisartan. To the best of our knowledge, no prior reports of telmisartan exist with respect to this direct effect on insulin secretion.

Moreover, our results showed that telmisartan, enhances insulin secretion in a glucose-dependent manner. Even at high concentrations (50 μM), no insulinotropic effect of telmisartan was observed under low glucose conditions (2.8 mM) (Fig. 1A). This indicated that telmisartan might be applied as an insulin secretagogue without the risk of hypoglycemia. Hypoglycemia is a frequent and severe adverse effect. Not only can apparent hypoglycemia cause coma or the disruption of daily life, but unrecognized, recurrent hypoglycemia can also lead to life-threatening cardiac complications such as arrhythmias and myocardial ischemia, and cause permanent cognitive impairment that may accelerate the onset of dementia*(Frier, 2014)*. In addition, emerging evidence suggests that forcing the β-cells to secrete insulin constantly, termed insulin hypersecretion, might have the potential to accelerate the decline in β-cell function and thus may constitute a contributing factor to the progression of T2DM *(Rustenbeck et al., 2010; Aston-Mourney et al., 2008)*. Therefore, glucose-independent insulinotropic agents have exhibited poor durability in maintaining long-term glycemic control *(Kahn et al., 2006)*. In comparison, our study showed that telmisartan increased insulin secretion in a manner proportional to the accumulating glucose concentration, thereby avoiding the risk of overstimulating the β-cells.

By means of its function as both an ARB and a partial agonist for PPAR-γ, telmisartan provides numerous beneficial effects in ameliorating T2DM and related complications*(Makino et al., 2008; Li et al., 2012; Nagel et al., 2006; Hasegawa et al., 2009; Saitoh et al., 2009; Yamana et al., 2008; Perl et al., 2010; Goyal et al., 2008)*. However, our results demonstrated that telmisartan also functioned rapidly as an insulin secretagogue, consequent to its unique electrophysiological effects on ion channels, which were independent of the AT1 receptor and PPARγ.

Glucose-induced insulin secretion and increase of [Ca^2+^]_i_ are tightly controlled by ion channels that regulate cell membrane potential. The closure of ATP-sensitive potassium (K_ATP_) channels caused by high glucose results in membrane depolarization and opening of Kv channels and VGCCs *(Sabatini et al., 2019; Kalwat and Cobb, 2017)*. Kv channels mediate repolarization of β-cells, and antagonize the Ca^2+^ influx induced by VGCC activation. Blockade of Kv channels therefore prolongs action potential duration, leading to an increase of insulin secretion. In support of this notion, here we found that inhibition of Kv channels was linked to telmisartan-induced augmentation of GSIS.

Moreover, we identified that telmisartan directly inhibited Kv2.1 channel. The Kv2.1 channel, as a Kv family member, accounts for the majority of Kv currents on β-cells, serving to not only negatively regulate GSIS but also potentiate β-cell apoptosis *(Kim et al., 2012; Tingting et al., 2018)*. Previous studies attributed telmisartan-induced protective effects against β-cells apoptosis and dysfunction to its action on the AT1 receptor and PPARγ*(Li et al., 2012; Hasegawa et al., 2009; Saitoh et al., 2009; Wang et al., 2019)*, however, our results indicated that the inhibition of Kv2.1 might also be involved. Moreover, based on its dual effects including regulation of insulin secretion and β-cell apoptosis, Kv2.1 is considered as a promising therapeutic target for T2DM by most researchers in the field. However, despite the occasional reports of small molecule Kv2.1 inhibitor*(Tingting et al., 2018; Zhou et al., 2016)*, no specific drugs have been developed for therapeutic use. Alternatively, as drug repurposing has become a successful approach to accelerate novel anti-diabetic drug development*(Turner et al., 2016)*, our favorable finding provides insight with regard to new options for anti-diabetic drug discovery. Furthermore, as Kv2.1 also serves as the key channel during neuronal apoptosis and its cleavage inhibits neuronal apoptosis *(Liu et al., 2018; Yao et al., 2009)*, the potential neuroprotective role of telmisartan also warrants further investigation. It should be noted here that there are many isoforms of the Kv channel contributing to the regulation of GSIS in β-cells *(MacDonald P et al., 2001)*; Accordingly, our data did not exclude the possibility that telmisartan also interacts with other Kv channel isoforms.

Of note, although the potent Kv channel inhibitor TEA blocks the majority of Kv channels, we found that telmisartan showed a more effective potentiation on insulin secretion and [Ca^2+^]_i_ concentration in the presence of TEA. Indeed, our findings revealed that in addition to Kv channels, VGCCs mediated the effects of telmisartan, which were also independent of the AT1 receptor and PPARγ. Moreover, we concluded that K_ATP_ channels were unlikely to be involved in telmisartan -regulated insulin secretion for several reasons. Specifically, the insulinotropic effect of inhibition of K_ATP_ channels is glucose-independent *(Dukes et al., 1994; Henquin, 2011)*, whereas telmisartan did not enhance insulin secretion under low glucose (2.8 mM) conditions (Fig. 1A and D). Conversely, the K_ATP_ antagonist tolbutamide increased [Ca^2+^]_i_ concentrations in β-cells under low glucose conditions (Fig. 2 A and B), suggesting that telmisartan and tolbutamide act on separate targets.

In summary, our results showed that beyond AT1 receptor blockade or PPARγ activation, telmisartan also inhibits Kv channels and activates VGCCs to promote extracellular Ca^2+^ influx, thereby enhancing [Ca^2+^]_i_ levels and amplifying GSIS. Our findings provide a new understanding of an anti-diabetes mechanism for telmisartan that is distinct from that of other ARBs, and may have important implications for determination of the choice of ARBs for the treatment of patients with both hypertension and diabetes. In addition, our identification of telmisartan also acting as a Kv2.1 inhibitor and glucose-dependent insulinotropic agent, provides a foundation for the development of new anti-diabetic drugs.

## Methods

### Animals

Adult male Wistar rats, weighing 240–260 g, were purchased from Beijing Weitong Lihua experimental animal center (Beijing, PR China). Five-week-old male diabetic *db/db* mice (BKS - Lepr^em2Cd479^/Gpt, stock number T002407) were obtained from GemPharmatech Co.,Ltd (Nanjing, China). Rats and mice were maintained in specific-pathogen-free surroundings, with a 12 h-light/dark cycles under controlled temperature (22 ± 2°C) and humidity (55–60%) conditions, and with free access to water and food. All animal care and experimental procedures conformed to the ethical guidelines for animal research at Shanxi Medical University and were approved by the Animal Care and Use Committee of Shanxi Medical University (Taiyuan, China).

### culture of islets and cells

The rat pancreas was isolated following injection of 1 mg/mL collagenase P (Roche, Indianapolis, IN, USA) through the common bile duct. After digestion at 37 °C for 11 min and density gradient centrifugation with Histopaque-1077 (Sigma-Aldrich, St.Louis, MO, USA) for 23 min, the expanded pancreas was dispersed, and islets remaining in the supernatant separated from the sediment. The islets were hand-collected under a dissection microscope, and single islet cells were obtained from islets using Dispase II (Roche, Indianapolis, USA) digestion. The *db/db* mouse islets were similarly obtained, although the pancreas was injected with 1 mg/mL collagenase V (Roche, Indianapolis, USA), then digested for 16 min and centrifuged twice with Hanks Balanced Salt Solution. Isolated islets and cells were cultured in RPMI 1640 (Hyclone, Thermo Scientific, Waltham, MA, USA) medium containing 11.1 mM glucose, supplemented with 10% fetal bovine serum, 1% penicillin and streptomycin at 37 °C in a humidified atmosphere of 5% CO2, 95% air.

Chinese hamster ovary (CHO) cells were obtained from the National Infrastructure of Cell Line Resource (Beijing, China). Lentivirus vectors overexpressing voltage-dependent potassium (Kv) 2.1 channels were constructed (Shanghai Genechem Co., Ltd., Shanghai, China) to transfect CHO cells, and the CHO-Kv2.1 cell line was established. CHO-Kv2.1 cells were cultured in Dulbecco’s modified Eagle’s medium (Hyclone, Thermo Scientific, Waltham, MA, USA) containing 4500 mg/L glucose in addition to 10% fetal bovine serum, 1% penicillin and streptomycin and 0.5 μg/mL puromycin (Beijing Solarbio Science & Technology Co., Ltd., Beijing, China). CHO cells were cultured under similar conditions except for puromycin selection.

### Insulin secretion assay

Handpicked separated islets were cultured for 1–2 days before the experiment. A total of five islets per tube were pre-incubated in Krebs Ringer bicarbonate-HEPES (KRBH) buffer under 2.8 mM glucose conditions for 30 min. The KRBH buffer contained 128.8 mM NaCl, 4.8 mM KCl, 1.2 mM KH2PO_4_, 1.2 mM MgSO_4_, 2.5 mM CaCl_2_, 5 mM NaHCO_3_, and 10 mM HEPES, adjusted to pH 7.4 with NaOH prior to the addition of 2% bovine serum albumin. Islets were then treated with different drugs and glucose conditions as indicated, and supernatant liquid was collected at the end of every 30 min incubation, and stored at −20 °C for insulin concentration measurement. Insulin secretion was determined using an Iodine [^125^I] Insulin Radioimmunoassay Kit (North Biological Technology Research Institute of Beijing).

### Calcium imaging technology

Calcium imaging was carried out at 28–30 °C using the calcium-sensitive dye Fura 2-AM (Dojindo Laboratories, Kumamoto, Japan), using an OLYMPUS IX71 inverted microscope and Meta Fluor software 7.8 (Molecular Devices, Sunnyvale, CA, USA). Islet cells were cultured on coverslips coated with adhesion reagent for 6–10 hour, then were loaded with 2 μM Fura 2-AM in KRBH buffer with addition of 2.8 mM glucose for 30 min at 37 °C. Subsequently, the loading buffer was removed, and cells were washed twice with KRBH solution to remove excessive fluorescent dye. Fura-2 was excited at 340 and 380 nm wavelengths in 1 s intervals with fluorescence emission detected at 510–520 nm wavelengths. The ratio of fluorescence intensity (F340/F380) was recorded to measure intracellular Ca^2+^ concentrations.

Fura 2-AM-loaded islet cells on coverslips were transferred to a glass chamber containing KRBH buffer with appropriate glucose conditions. Between each test, the reagent was dripped onto the coverslip and F340/F380 data points were acquired to monitor the changes of intracellular Ca^2+^ level. The average value during 30 s F340/F380 spikes (15 s before and after the peak of F340/F380) for each test was used to compare the change of Ca^2+^ concentrations under different treatments, unless otherwise stated.

### Electrophysiology

Whole-cell recording patch-clamp technology was applied to detect voltage-activated currents and record action potentials using an EPC-10 amplifier and PULSE software from HEKA Electronik (Lambrecht, Germany) at room temperature. Islet cells were cultured on glass coverslips coated with cell adherent reagent (Applygen Technologies Inc., Beijing, China).

In voltage-clamp mode, to record Kv currents, patch pipettes (5–8MΩ) were loaded with intracellular solution containing 10 mM NaCl, 1 mM MgCl_2_, 0.05 mM EGTA, 140 mM KCl, 0.3 mM Mg-ATP, and 10 mM HEPES, pH 7.25 adjusted with KOH. Cells were transferred to a recording chamber containing extracellular solution consisting of 138 mM NaCl, 5.6 mM KCl, 1.2 mM MgCl_2_·6H_2_O, 2.6 mM CaCl_2_, 11.1 mM glucose, and 5 mM HEPES (pH 7.4 adjusted with NaOH). The β-cells were identified by cell capacitance (>7 pF) *(Göpel et al., 1999)*and were clamped to a holding potential of −70 mV, then test potentials were elicited by ranging from −70 mV to 80 mV in 10 mV steps for 400 ms.

For voltage-gated Ca^2+^ channel (VGCC) currents, the intracellular solution contained: 120 mM CsCl, 20 mM TEA (Sigma-Aldrich), 5 mM MgATP, 1 mM MgCl_2_, 0.05 mM EGTA, and 10 mM HEPES (pH 7.25 adjusted with CsOH). The extracellular solution consisted of: 100 mM NaCl, 20 mM TEA, 20 mMBaCl_2_, 4 mM CsCl, 1 mM MgCl_2_, 5 mM HEPES, and 3 mM glucose (pH 7.4 adjusted with NaOH). Ca^2+^ was replaced with Ba^2+^ as the charge carrier in the extracellular solution to eliminate Ca^2+^- dependent inactivation of the VGCCs. β-cells were clamped to a holding potential of −70 mV, and then elicited by test potentials of −50 mV to 30 mV in 10 mV steps for 50 ms.

In current-clamp mode, β-cells were elicited by 4 ms currents of 150 pA to record action potentials. The time between the initiation and the point where membrane potential returned to within 10 mV of the resting membrane potential, was considered to be the measurement of action potential duration.

### In vivo evaluation of mice and drug administration

At the age of 8 weeks, the mice were given fasting glucose teste to ensure that diabetes models were successfully established. Given that a therapeutic doses of telmisartan are 40–80 mg/day in humans, the conversion for mice was approximately 8.2–16.4 mg/kg of body weight *(Nair and Jacob, 2016)*. In our experiment, the mice were administered acute oral acute oral telmisartan treatment at 15 mg/kg of body weight. At 2 hours following drug intake, when the onset of action of telmisartan reached a maximum, oral glucose tolerance test (OGTT) was performed *(Gohlke et al., 2001)*.

### OGTT

At the age of 8 weeks, following overnight fasting (14 h), the mice were randomly divided into groups receiving treatment with telmisartan (in drinking water containing 0.5% carboxymethyl cellulose sodium salt) or vehicle by gavage. For OGTTs, groups of mice were fed with glucose at 1.5 g/kg body weight orally, then a blood sample was collected from the tail vein and glucose levels were assessed using a Sinocare Glucometer (Changsha, China) at baseline (0 minute) and after 15, 30, 60, 90, and 120 min. At the age of 11 weeks, the mice were treated as described above and additional blood samples (50 μL) were obtained in a heparinized microhematocrit tube at 0, 15, 30, 60, 90, and 120 min. After centrifugation, the plasma was collected for insulin concentration measurement using the Mercodia Mouse Insulin ELISA (stock number 10-1247-01, Uppsala, Sweden).

### Statistical analysis

All experimental data are presented as the means ± SEM. P< 0.05 was considered to indicate statistical significance. Sharpiro–Wilk tests were used to analyze the normality of the data. Upon normal distribution, the means of numerical variables were compared using the Student’s *t* test or one-way analysis of variance (ANOVA), whereas data with non-normal distribution were analyzed using the Mann–Whitney Rank Sum Test or Kruskal–Wallis one-way ANOVA on Ranks. If any statistically significant difference was detected among three or more groups, the Student–Newman–Keuls method or Tukey test was performed for post hoc comparisons, unless otherwise stated.

## Acknowledgements

This work was supported by NSFC (81670710; 81770776; 81973378), 136 project in Shanxi Bethune Hospital(2019XY015), Cultivate Scientific Research Excellence Programs of Higher Education Institutions in Shanxi (2019KJ022).This research project was supported by Shanxi Scholarship Council of China (2017-053), FSKSC and 1331KSC, Department of Education Innovation Project in Shanxi Province (2019BY078), and Shanxi Youth Science and Technology Research Fund (201901D211323).

## Author contributions

Y. Z., Y. L. and T. L. conceived and designed the study; T. L. performed the ex vivo experiments with assistance from L. C., H. X., X. Y., M .L., T. B., Z. L., and Q.G.; T. L., L. C., H. Y., and L. Z. carried out the in vivo experiments; T. L., M. Z., and P. H. analyzed the data. T. L., Y. Z., and Y. L. wrote the manuscript.

---

**Fig. S1:**
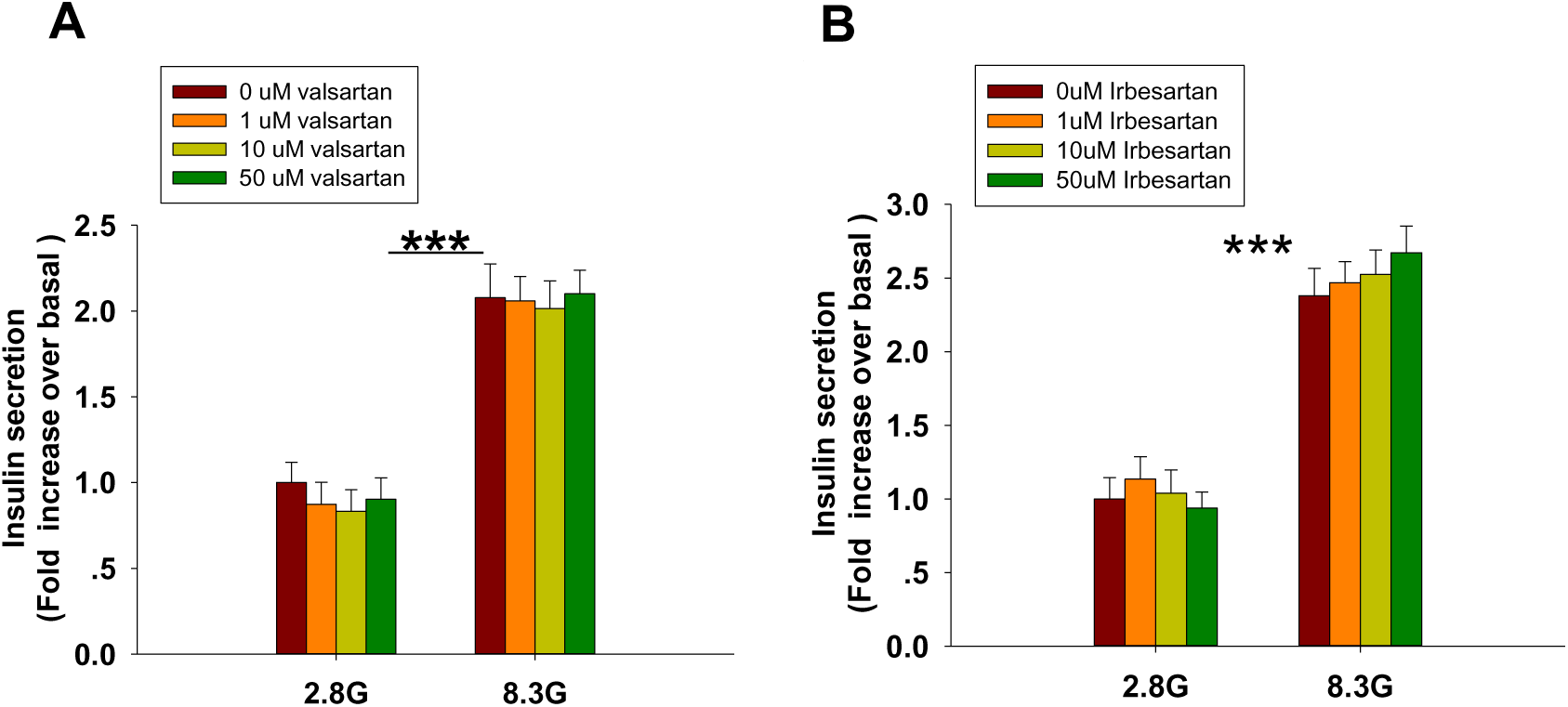
Neither valsartan nor irbesartan potentiated insulin secretion in rat islets (n = 7 tubes per group). (A) Rat islets were treated with various doses (1, 10, and 50 μM) of valsartan under 2.8 mM and 8.3 mM glucose (denoted as 2.8 G and 8.3 G) conditions. (B) Rat islets were treated with various doses (1, 10, and 50 μM) of irbesartan under 2.8 mM and 8.3 mM glucose conditions. All results are normalized to basal secretion at 2.8G, and reported as the means ± SEM. Statistical differences among groups were compared using one-way analysis of variance (ANOVA) and Student–Newman–Keuls method post hoc analysis. *** P < 0.001.

**Fig. S2:**
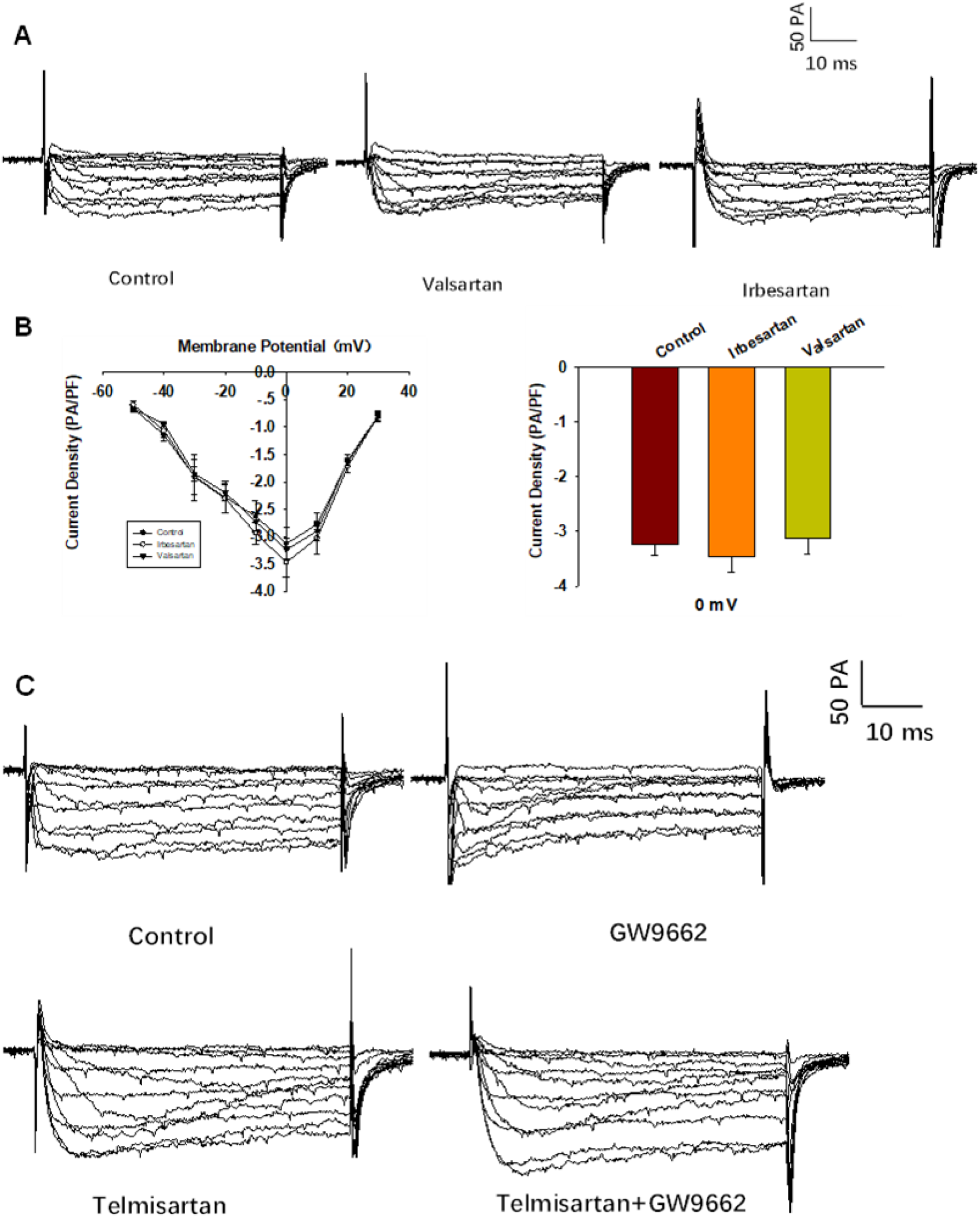
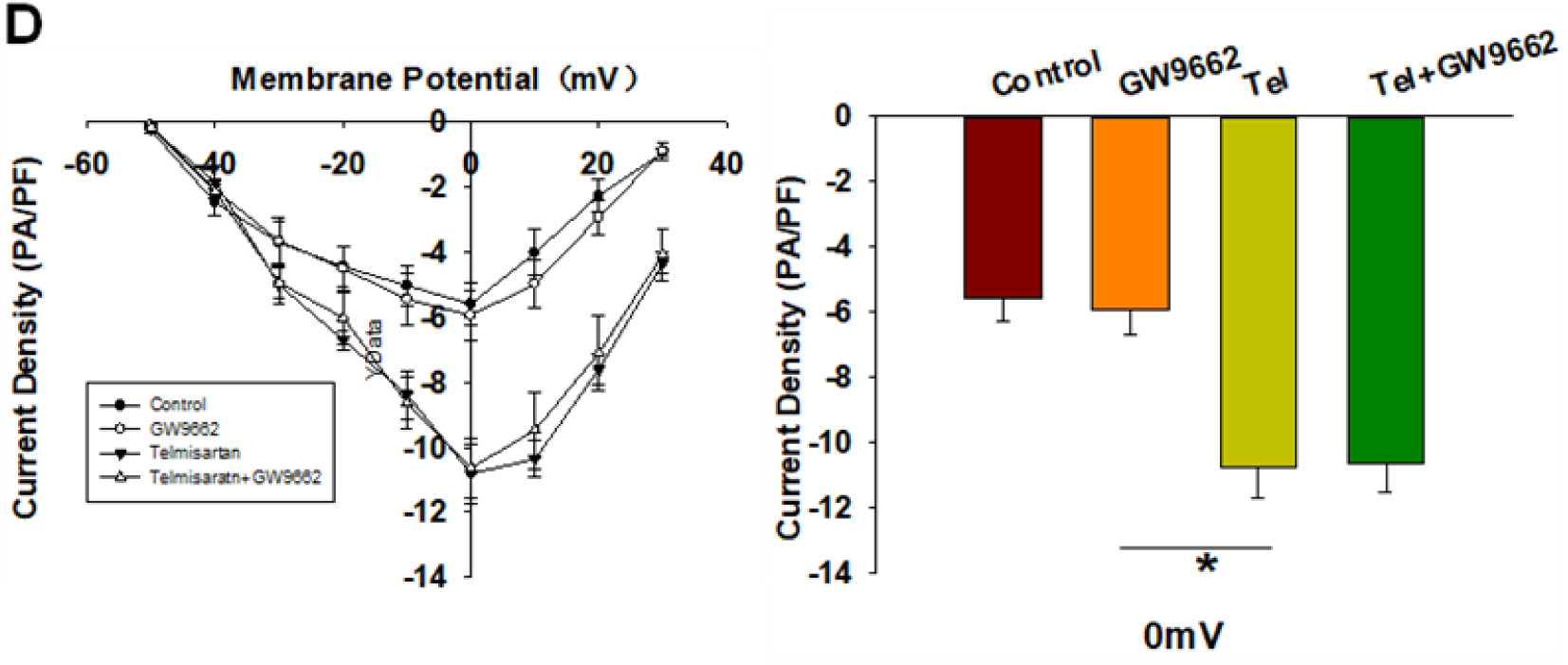
The AT-1 receptor and PPARγ are not involved in telmisartan-induced activation of VGCCs. (A) Representative current traces recorded with treatment of valsartan (10 μM) and irbesartan (10 μM) in β-cells. (B) Current-voltage relationship curves and the summary of the mean Ca^2+^current density recorded at 0 mV depolarization (n = 7). (C) Representative current trances recorded under treatment of telmisartan (10 μM) alone or in combination with GW9662 (10 μM) in β-cells. (D) Current-voltage relationship curves and the summary of the mean Ca^2+^ current density recorded at 0 mV depolarization (control, n = 10; GW9662, n = 6; telmisartan, n = 8; telmisartan+GW9662, n = 6). All results are reported as the means ± SEM. Statistical differences among three or more groups were compared using one-way ANOVA. For comparing the effects of GW9662 groups, Dunn’s method post hoc analysis was applied. *P < 0.05

**Fig. S3:**
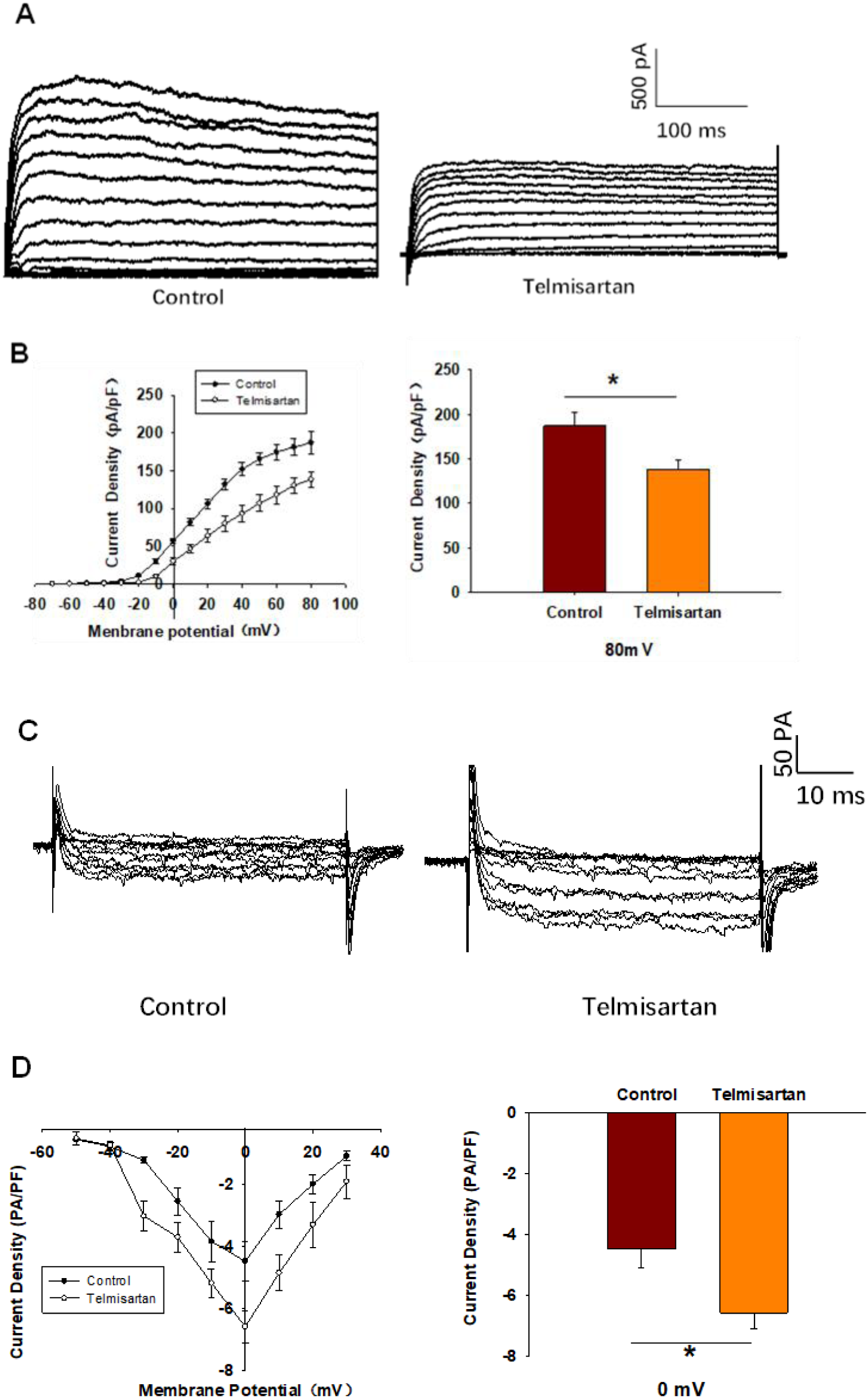
Telmisartan exerts similar electrophysiological effects on β-cells of db/db mice. (A) Representative Kv channels current trances recorded with treatment of telmisartan (10 μM) in β-cells. (B) Current-voltage relationship curves and the summary of the mean current density of Kv channels recorded at 80 mV depolarization (n = 6). (C) Representative Ca^2+^ current trances recorded with treatment of telmisartan (10 μM) in β-cells. (D) Current-voltage relationship curves and the summary of the mean Ca^2+^ current density recorded at 0 mV depolarization (control, n = 6; telmisartan, n = 7). All results are reported as the means ± SEM. Statistical differences between two groups were determined using an unpaired two-tailed Student’s *t* test. *P< 0.05.

